# Dielectrophoresis assisted rapid, selective and single cell detection of antibiotic resistant bacteria with G-FETs

**DOI:** 10.1101/842187

**Authors:** Narendra Kumar, Wenjian Wang, Juan C. Ortiz-Marquez, Matthew Catalano, Mason Gray, Nadia Biglari, Kitadai Hikari, Xi Ling, Jianmin Gao, Tim van Opijnen, Kenneth S Burch

**Affiliations:** Department of Physics, Boston College, Chestnut Hill, Massachusetts 02467, United States; Department of Biology, Boston College, Chestnut Hill, Massachusetts 02467, United States; Department of Chemistry, Boston College, Chestnut Hill, Massachusetts 02467, United States; Department of Chemistry, Boston University, Boston, Massachusetts 02215, United States; Division of materials science and engineering, Boston University, Boston, MA, 02214, United States; The photonics center, Boston University, Boston, MA, 02214, United States

**Keywords:** Biosensors, G-FET, Peptide probes, Dielectrophoresis, Antibiotic resistance bacteria, Electrical detection

## Abstract

The rapid increase in antibiotic resistant pathogenic bacteria has become a global threat, which besides the development of new drugs, requires rapid, cheap, scalable, and accurate diagnostics. Label free biosensors relying on electrochemical, mechanical, and mass based detection of whole bacterial cells have attempted to meet these requirements. However, the trade-off between selectivity and sensitivity of such sensors remains a key challenge. In particular, point-of-care diagnostics that are able to reduce and/or prevent unneeded antibiotic prescriptions require highly specific probes with sensitive and accurate transducers that can be miniaturized and multiplexed, and that are easy to operate and cheap. Towards achieving this goal, we present a number of advances in the use of graphene field effect transistors (G-FET) including the first use of peptide probes to electrically detect antibiotic resistant bacteria in a highly specific manner. In addition, we dramatically reduce the needed concentration for detection by employing dielectrophoresis for the first time in a G-FET, allowing us to monitor changes in the Dirac point due to individual bacterial cells. Specifically, we realized rapid binding of bacterial cells to a G-FET by electrical field guiding to the device to realize an overall 3 order of magnitude decrease in cell-concentration enabling a single-cell detection limit, and 9-fold reduction in needed time to 5 minutes. Utilizing our new biosensor and procedures, we demonstrate the first selective, electrical detection of the pathogenic bacterial species *Staphylococcus aureus* and antibiotic resistant *Acinetobacter baumannii* on a single platform.

## 1. Introduction

The over prescription and misuse of antibiotics is causing a surge in the number of antibiotic resistant bacterial infections around the world(Centres for 2013; López-Góngora et al. 2015; Scholte et al. 2015) Central to solving this crisis are cheap, rapid and easy diagnostic methods that accurately identify the bacterium causing the infection and its associated antibiotic resistance profile. Antibiotic susceptibility testing (AST) is mostly carried out by phenotypic methods that require prior identification of bacterial pathogens from patients (at the species and/or strain level) and incubation under antibiotic conditions,(Syal et al. 2017; Varadi et al. 2017) a lengthy process that can take up to 24 h to a month depending on the species.(Jenkins and Schuetz 2012) Moreover, both species/strain identification and AST require trained specialists, specific laboratory environments and often expensive instrumentation.(Syal et al. 2017; van Belkum et al. 2019; Varadi et al. 2017) Since these conditions limit widespread application and implementation into actual treatment strategies at most points-of-care, there is much room for improvement to develop new diagnostic devices that have the potential for adoption across a large variety of use cases. Ideally these devices would be cheap, easy to implement, scalable, and accurately identify both the pathogen as well as its antibiotic resistance profile with high specificity and sensitivity.(Boucher et al. 2016; Huang et al. 2015; Zumla et al. 2014)

Label free biosensors such as electrochemical, mechanical, and mass based could meet these requirements and as such have been widely studied in the detection of whole bacterial cells. (Ahmed et al. 2014; Templier et al. 2016) However, the trade-off between selectivity and sensitivity of such sensors remains a key challenge as they rely on highly specific probes combined with a sufficiently sensitive and accurate transducer that can be miniaturized and multiplexed and that is cheap and easy to use. Knowing that bacteria possess negative surface charge at physiological pH, (Kłodzińska et al. 2010; Soon et al. 2010) graphene field effect transistors (G-FETs) can be an excellent choice as they have high sensitivity in detection of proteins and DNA, scalability, biocompatibility and ease of incorporation on conventional and flexible substrates.(Fu et al. 2017a; Fu et al. 2017b; Ping et al. 2016; Xu et al. 2017) An impediment to the broad use of G-FETs for bacterial sensing lies in the lack of suitable probes which should be readily available, easily handled (simple preparation and/or long shelf life), and have species, strain, and resistance-specificity.(Rubab et al. 2018; Templier et al. 2016) An equally crucial challenge is enhancing the sensitivity to achieve detection at a clinically relevant cell density.

Progress in this field has been limited, in part, by the typical large charge induced by the attachment process or the probes themselves, reducing sensitivity to the additional charge of the target. Additionally, if the probes are too big, the target can be beyond the Debye screening length of the solution, resulting in reduced sensitivity.(Donnelly et al. 2018) These factors remain the reason behind the limited utilization of G-FETs for detection of bacteria, resulting in only a single report for a lab strain of *Escherichia coli.*(Huang et al. 2011; Wu et al. 2017) Therefore, smaller size, neutral nature and high selectivity would be the desired features of optimal probes for utilization in G-FET based biosensors. Peptide probes provide an advantage over antibodies or aptamers, due to their small size, neutral nature (no net charge), long stability, and easy synthesis (Mannoor et al. 2012; Mannoor et al. 2010; McGregor 2008; Zasloff 2002). Additionally, synthetic peptides obtained after probe screening using phage display library (Liu et al. 2018b; Liu et al. 2016a) in a colorimetric immunoassay prove to be relatively inexpensive to produce and solve the issue of lower level of selectivity for instance observed with antimicrobial peptides.(Etayash et al. 2014; Liu et al. 2016b; Mannoor et al. 2012; Mannoor et al. 2010; Soares et al. 2004) Similarly, a previous work by some of us developed a phage display platform that can rapidly select for small peptides that recognize and bind specific bacterial species or strains. These chemically modified peptide probes showed submicromolar affinity and high specificity for clinical strains of *Staphylococcus aureus* and *Acinetobacter baumannii.* (McCarthy et al. 2018)

There are no reports on the sensing of clinically relevant pathogenic bacteria and antibiotic resistant strains utilizing G-FETs. Most studies utilizing G-FETs have relied on changes in source-drain current, due to relatively small shifts in Dirac voltage (*V_D_*) upon attachment of their target. Beyond limiting detection limits, this approach requires sensing small changes in voltage that are more sensitive to noise. This results from the fixed amount of charge per target, whereas the shift is dependent on the induced charge density.(Wu et al. 2017) As such the effect of the bacteria on a single G-FET could be enhanced with smaller active areas, however this also requires a far higher cell density, potentially at a level that is much above what is considered to be clinically relevant. Nonetheless, enabling further reduction of the needed active area, would also allow for multiplexing by placing numerous G-FET elements with a variety of probes in the sample space.

Herein, we provide a solution to these problems by implementing for the first time in a G-FET device, specific synthetic peptides as probes and dielectrophoresis realized by electric field assisted binding to obtain a clinically relevant detection limit. This enabled the use of small active area G-FETs (10 × 40 μm) producing single cell resolution, wherein large changes in *V_D_* are directly observed by measuring the device resistance versus gate voltage. Additionally, we obtained high specificity to targeted pathogenic bacteria including a Gram positive *S. aureus* strain and a Gram negative antibiotic resistant *A. baumannii* strain. The G-FETs design, with two independent wells (with 3 devices in each well) on a single 1×1 cm chip, required only 20 μl of sample. Utilizing this design, we can simultaneously detect two different bacterial species, or test specific and unspecific peptide probes on the same platform. Further, the electric field assisted binding of these bacteria resulted in a lowering of the limit of detection to 10^4^ cells/ml, which is within a clinically relevant regime, and a reduction in the time of detection to 5 minutes.

## 2. Material and methods

### 2.1. G-FET Fabrication and Characterization

G-FETs were fabricated on chemical vapor deposition (CVD) monolayer graphene transferred over SiO_2_/Si substrates. Monolayer graphene was grown on a copper foil via low pressure CVD. The copper foil (Alfa Aesar) was pre-treated in Ni etchant (Transene) to remove any coatings or oxide layers from its surface. The tube furnace was evacuated to a read pressure of 200 mTorr with a constant flow of H_2_ (10 sccm). Prior to growth, the foil was annealed at 1010 °C (ramp rate 25 °C/min) for 35 minutes. Growth was done at 1010 °C with 68 sccm of H_2_ and 3.5 sccm of CH_4_ for 15 minutes. After growth, a polymethyl methacrylate (PMMA) layer was spin coated on one side of the copper foil and baked for 60 seconds at 60 °C. To facilitate smooth and fast etching of the copper foil, the backside graphene was etched out using oxygen plasma with 60 watt power for 60 seconds. The exposed copper was etched away in Ni etchant for 2h at 60 °C. The remaining PMMA/graphene structure was washed in 2 water baths, the first water bath for 60 seconds and the second for 30 minutes, to rinse away left over etchant. The PMMA/graphene was transferred onto a SiO_2_/Si chips of size 1 × 1 cm. Any leftover water was slowly dried out with nitrogen gas, and finally the PMMA was dissolved in acetone vapors; isopropanol alcohol (IPA) (Fischer) was used for a final wash. The chips were baked at 300 °C for 8h in vacuum followed by deposition of 3 nm AlOx at room temperature by evaporating aluminum at oxygen pressure of 7.5 × 10^5^ mbar. Substrates were baked at 175 °C for 10 minutes before lithography process. The electrodes patterning was done using bilayer photoresist (LOR1A/S1805) and laser mask writer (Heidelberg Instruments) followed by Au/Cr (45 nm/5 nm) deposition and lift off using remover PG (MicroChem). After that the graphene patterning was done with lithography using same bilayer resist and oxygen plasma etching. Devices were cleaned with remover PG and rinsed with IPA, DI water and dried with Argon. In order to protect the electrodes and edges of the graphene for liquid gating, photolithography was done using S1805 to open the sensing area (10 × 40 μm) and contact pads while leaving remaining chip covered. The developing time was increased to 90 seconds to etch away the AlOx layer deposited in the beginning to protect the graphene from photoresist. Finally, the chips were baked at 150 °C for 5 minutes and then temperature increased to 200 °C and baked for 5 more minutes to harden the photoresist. To hold the solution for the measurement, two PDMS wells of size 2.5 × 2.5 mm were fabricated and placed over the chip having two sets of the devices with three devices in each well.

### 2.2. Peptide synthesis

Solid phase peptide synthesis was performed on a rink amide resin using Fmoc chemistry. An alloc-protected diaminopropionic acid residue was installed at the C-terminus for on-resin coupling of pyrene. The alloc protecting group was selectively removed by tetrakis (triphenylphosphine) palladium (0) and phenylsilane in DCM for 1 h. 2 equivalents of 1-Pyrenebutyric acid N-hydroxysuccinimide ester in 20% v/v DIPEA/DMF was added. The coupling was done in 2 h at room temperature. The peptides were cleaved off resin and globally deprotected with 90% TFA, 5% H_2_O, 2.5% triisopropylsilane, 2.5% 1,2-ethanedithiol for 2 h. Crude peptides were obtained via cold ether precipitation and purified by RP-HPLC. For cysteine alkylation, the peptides were treated with 3 equivalents of APBA-IA or IA in 5% v/v DIPEA/DMF for 3 h and purified via RP-HPLC. All peptides were characterized with LC-MS to confirm their identities and excellent purities (>95%).

### 2.3. Bacterial strains and culture conditions

Detections were made using the following strains: *S. aureus* (ATCC 6538), wild-type *A. baumannii* (AB5075),(Jacobs et al. 2014) colistin resistant and LOS deficient *A. baumannii* (5075 LOS−),(Boll et al. 2016) *B. subtilis*, and *E. coli (BL 21)*. All bacteria were cultured overnight in LB broth at 37 °C with 220 rpm constant shaking. The overnight culture was diluted 10^2^ times in fresh media and grown to an OD_600_ of 0.5-1.0. These fresh cultures were then washed and diluted with 1x PBS (pH 7.4) buffer to obtain the desired concentrations.

### 2.4. G-FET functionalization and measurement conditions

G-FETs were functionalized with peptides by incubating with 10 μM concentration of P-KAM5_Probe and P-KAM8_Probe for optimized durations of 2 h and 16 h, respectively (Figure S1). The method used to find these times is outlined in the supplemental. To minimize the noise in the electrical measurement, G-FETs were characterized by measuring the resistance using a digital multimeter by sweeping Liquid gate voltage between 0 V to a maximum of 1.7 V. The test current was limited to 10 μA reduce the effects of heating the device and prevent failure. The maximum (Dirac voltage) in resistance versus voltage plot was chosen as reference point and shift in the maximum was measured upon bacterial binding. Functionalized G-FETs were incubated with 20 μl desired bacterial solutions in 1x PBS (pH-7.4), while the measurements were performed in 0.01x PBS diluted in DI water to maximize the signal by reducing the Debye screening effect (Figure S2).(Formisano et al. 2016; Wu et al. 2017). A platinum wire of 0.5 mm diameter was used for liquid gating.

## 3. Results and discussion

### 3.1. G-FET device construction and baseline measurements

G-FET devices consist of a low pressure chemical vapor deposition (CVD) graphene on a standard SiO_2_/Si substrate, and etched into an active area of 20 × 50 μm, with Cr/Au source and drain. The contacts were passivated and the sensing area (10 × 40 μm) was defined easily with a conventional hard baked photoresist (S1805), instead of a dielectric (Figure 1a). To measure the baseline conductance/resistance of a device, they were tested in liquid gate mode where a Pt wire was chosen as reference electrode and 0.01x PBS as electrolyte (Figure 1b). The measured average Dirac voltage (*V_D_*) of the fabricated G-FETs is around 0.7±0.16 V, consistent with the surface potential of the platinum wire and diluted concentration of PBS.(Chen et al. 2013) The observed variation in the *V_D_* with devices made on various substrates and batches is attributed to the impurities at the graphene/SiO_2_ interface induced during the graphene transfer process. The mobility was calculated by linearly fitting the hole and electron regimes of conductance (σ) versus voltage (V_G_) using 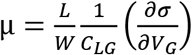, where L, W are the length and width of the channel, C_LG_ is the liquid gate capacitance. Value of C_LG_ was taken to be 1.65 μF/cm^2^ based on the sum of the quantum capacitance (C_Q_) of graphene and electric double layer capacitance (C_DL_) consistent with 0.01x PBS.(Wu et al. 2017) Results from a high mobility device are shown in Figure 1b, while the average hole and electron mobility values obtained from different devices are ~670 ± 125 and ~690 ± 83 cm^2^/V·s, respectively. These mobilities are consistent with reported values for CVD graphene on SiO_2_ substrates.(Kireev et al. 2017; Wu et al. 2017)

**Figure 1.**
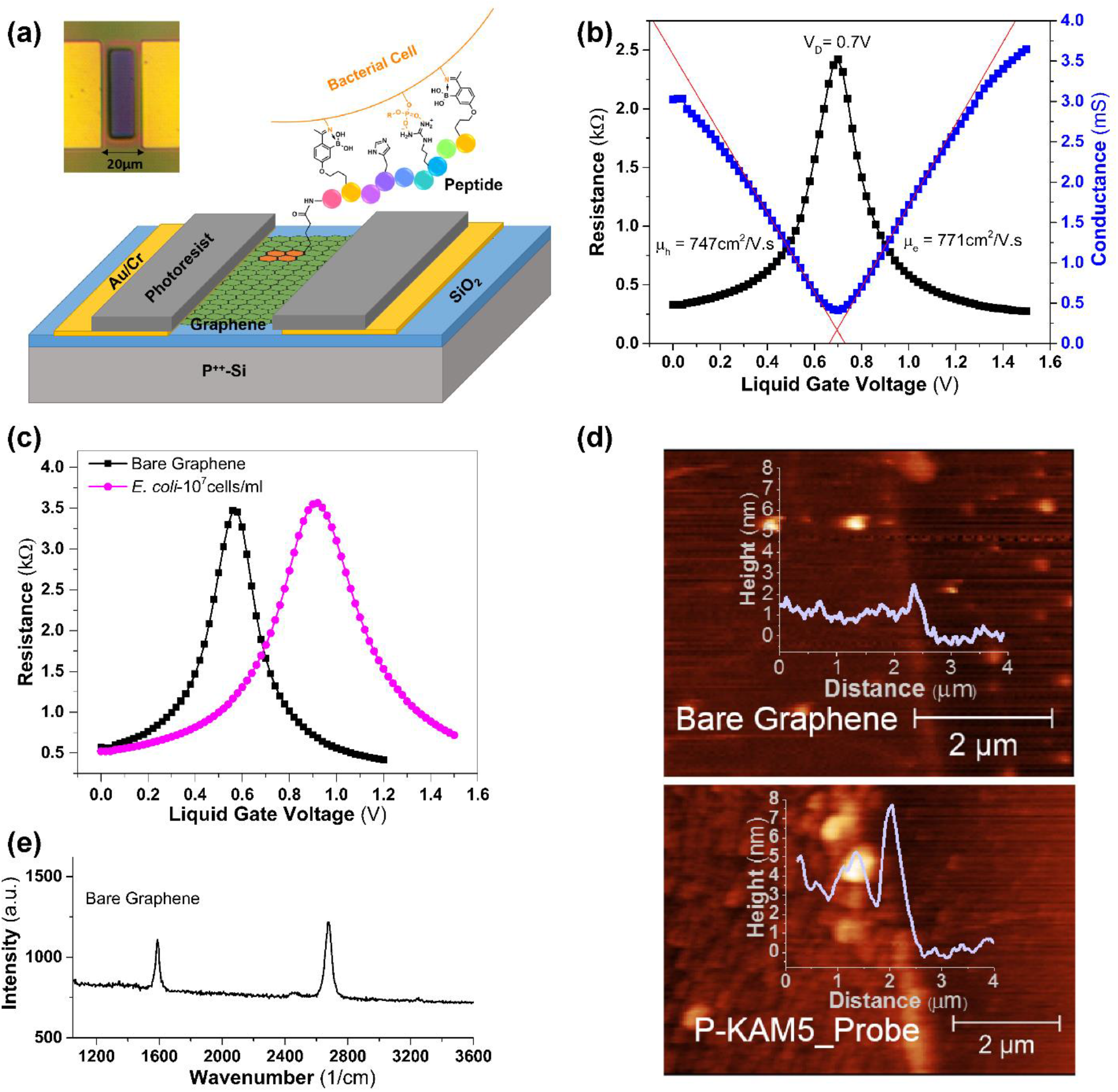
Scheme of functionalization and characterization of G-FETs. (a) A schematic of a G-FET functionalized with a pyrene-conjugated peptide probe binding to the surface of a bacterium. The inset shows a light microscopy image of G-FET, with an active area of 10 × 40 μm, located in between two gold contacts. (b) Resistance/conductance vs voltage plots of G-FET representing the Dirac voltage (0.7 V), hole and electron mobilities of 747 and 771 cm^2^/V.s. (c) G-FET characteristics before and after adsorption of E. coli. A shift of 360 mV in the Dirac voltage observed when G-FET with bare graphene was incubated with E. coli (pink circles) in comparison with the bare graphene (black squares). (d) AFM image of the patterned graphene before and after peptide functionalization. After functionalization, the coverage of the graphene channel by the P-KAM5_Probe peptide is shown as an increase in height of ~2.5 nm (lower panel) compared with the bare graphene surface (upper panel). (e) Raman spectrum (532 nm excitation) of CVD graphene transferred over SiO_2_/Si substrate. The spectrum shows 2D peak at 2,679 cm^−1^ and G peak at 1,587 cm^−1^ with I2D/IG ~1.6, suggesting the graphene is monolayer. Absence of D peak ~1350 cm^−1^ indicating minimal defects in graphene.

### 3.2. Bacterial detection using G-FETs with bare graphene

To ensure the dimensions of the device and operating conditions would detect bacteria, we first exposed a bare G-FET containing a strip of non-functionalized bare graphene to two types of bacteria: *E. coli* and *S. aureus*. Each species was incubated for 45 minutes on different devices at a bacterial suspension of 10^7^ cells/ml. As evident from Figure 1c, a strong shift in *V_D_* of 360 mV results from exposure to *E. coli* and 220 mV from *S. aureus* (Figure S3). This positive shift by attachment of *E. coli* is consistent with that observed in back gate mode.(Mulyana et al. 2018) These results confirm that our bare graphene is highly sensitive to the bacterial surface charge but cannot distinguish between different bacterial species, strains or resistance state, indicating that G-FET’s require specific probes to be integrated on such devices.

### 3.3. Peptide-pyrene conjugates enable a simplified single-step graphene functionalization process

In order to maintain the electronic properties of graphene, it is preferable to use probes with non-covalent functionalization through π-π stacking of pyrene-based linker molecules.(Georgakilas et al. 2012) This functionalization typically requires multiple steps, starting at linker attachment and followed by incubation with biosensing probes. As a result, G-FETs are exposed to different solvents with the potential of significantly affecting the doping level of the graphene.(Ping et al. 2016; Wang et al. 2015; Xu et al. 2017) This also makes the preparation and functionalization of devices complex, cumbersome, and potentially expensive. Previously we developed a phage display platform that can rapidly select for small peptides that recognize and bind specific bacterial species or strains.(McCarthy et al. 2018) KAM5 is one such peptide that was identified in our previous study to specifically detect *S. aureus*, showing an EC_50_ of ∼1.5 μM in a cell staining assay. To minimize exposure of G-FET to solvents and facilitate single step functionalization with the desired probe, we synthesized a peptide-pyrene conjugate (P-KAM5_Probe), by on-resin coupling of 1-pyrenebutyric acid N-hydroxysuccinimide ester (PBASE) onto the diaminopropionic acid residue installed at the C-terminus of the peptide. These pyrene-conjugated peptides are then simply dissolved in aqueous solution incubated on the device for 2 h followed by a wash step, with no additional chemicals or treatments required. In order to confirm uniform functionalization, P-KAM5_Probe was attached to the patterned bare graphene surface and characterized with atomic force microscopy (AFM; Figure 1d). The height of graphene functionalized with P-KAM5_Probe increased by ~2.5 nm as compared to the bare graphene surface. This height increase is expected and consistent with peptides attached to carbon nanotubes or graphene oxide,(Kuang et al. 2010; Liu et al. 2018a) confirming the attachment of the probe to the device. The peptide-pyrene conjugates thus facilitate simplified graphene functionalization by a single step process, which enables rapid and easy preparation of the device as well as reduced fabrication cost. Raman spectrum was carried out to confirm the quality of the CVD graphene used for the G-FET fabrication (Figure 1e). Obtained 2D peak at 2,679 cm^−1^ and G peak at 1,587 cm^−1^ with the ratio in their intensities i.e. I_2D_/I_G_ ~1.6, confirming the single layer graphene, while the absence of D peak ~1350 cm^−1^ indicating negligible amount of defects in graphene.(Wu et al. 2018)

### 3.4. Species specific detection of a Gram-positive bacterial pathogen

To test the potential of our G-FET design for detecting specific bacterial species, we functionalized G-FETs with P-KAM5_Probe, which is expected to bind *S. aureus*, or a control peptide (P-KAM5_Control, see Supporting Information for details), in which the key functional groups for binding are missing thereby making it incapable of binding bacteria. As shown in Figure 2a-b, functionalization of G-FET with either peptide does not show any shift in *V_D_*, consistent with their charge neutral structure at pH 7. Upon incubation with *S. aureus* at 10^7^ cells/ml on a GFET functionalized with P-KAM5_Probe a voltage shift of 300 mV was observed (Figure 2a). In contrast and as expected, no notable voltage shift was observed when *S. aureus* was incubated on devices functionalized with P-KAM5_Control (Figure 2b). To visually confirm that the shift in *V_D_* is due to bacteria attached to the surface of the graphene, devices were analyzed using optical microscopy. *S. aureus* is a spherically shaped bacterium with an approximate 1 μm diameter,(Monteiro et al. 2015) and black dots, representing individual bacterial cells, were observed only on devices functionalized with P-KAM5_Probe and not P-KAM5_Control (inset of Figure 2a-b). The observed positive shift in the *V_D_* is attributed to the negatively charged surface of bacterial cells which increase the hole carrier density in graphene.(Wu et al. 2017) To probe the postulated *S. aureus* specificity of our G-FET, the peptide functionalized devices were comparatively tested against *Bacillus subtilis* a different Gram-positive species and *E. coli* a representative Gram-negative bacterium. No significant shift in *V_D_* was observed when the devices were incubated with either species under the same conditions used for *S. aureus* (Figure 2c). Importantly, after rinsing with DI, the same devices were subsequently incubated with *S. aureus* resulting in an average shift in *V_D_* of ~190 mV, indicating that the devices functionalized with P-KAM5-probe are specific to *S. aureus* and insensitive to other Gram-positive and negative species.

**Figure 2.**
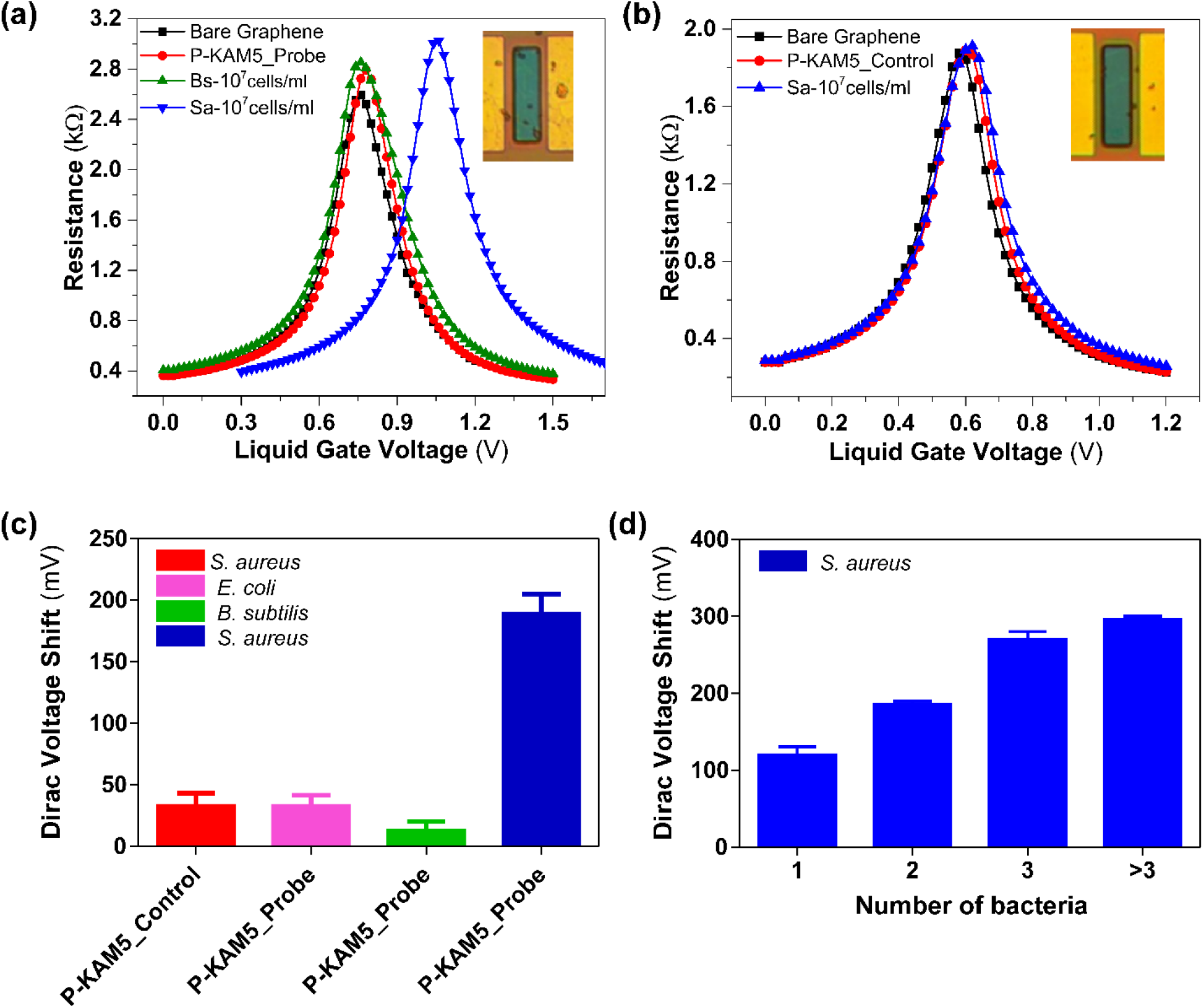
Specific detection results of S. aureus. Resistance vs voltage plots of G-FET for detection of S. aureus (a) G-FET functionalized with Probe peptides (P-KAM5_Probe) and incubated with B. subtilis and S. aureus at a concentration of 10^7^cells/ml. No shift was observed with peptides and B. subtilis while a shift of ~300 mV is seen with S. aureus as well as the attachment of bacterial cells to the graphene (see image in inset). (b) G-FET functionalized with control peptides (P-KAM5_Control) and incubated with S. aureus at a concentration of 10^7^ cells/ml. No voltage shift was observed after the incubations with peptide as well as with bacteria. Additionally, no attachment of bacterial cells on the graphene was observed (see image in inset). (c) Comparative values of average voltage shift with specific and unspecific detection of S. aureus. No notable shift was observed when G-FET were functionalized with P-KAM5_Control and incubated with S. aureus at a concentration of 10^7^ cells/ml. Furthermore, no notable shift was observed when G-FET were functionalized with P-KAM5_Probe and incubated with unspecific bacteria (E.coli and B. subtilis) at a concentration of 10^7^ cells/ml. An average shift of ~190 mV was observed when G-FET functionalized with P-KAM5_Probe and incubated with S. aureus at a concentration of 10^7^ cells/ml. (Data represents average and standard deviation of at least 6 independent replicates) (d) Measured Dirac voltage shift of G-FETs having different number of bacteria (S. aureus) attached. Devices having single bacterium attached show an average shift of ~130 mV and linear increase in voltage shift is observed with increased number of bacteria attached. (Data represents average and standard error of at least 3 independent replicates).

Additional correlation was found by using optical imaging to count the number of bacteria on each G-FET after electrical detection. Specifically, after measuring 20 devices functionalized with P-KAM5_Probe and incubated with *S. aureus* followed by inspection with light microscopy it was observed that a strong correlation exists between the number of bacteria that are bound by the probe to the graphene and the registered voltage shift. As shown in Figure 2d, a linear shift of *V_D_* was seen with increasing number of attached bacteria with a sensitivity of 56.3 ± 7.3 mV/bacteria. Surprisingly, the devices are able to detect the attachment of a single bacterium with an average voltage shift of 128±18 mV, a nearly 20% increase in the measured V_D_ over the as-prepared G-FET. Moreover, the voltage shifts of ~130 → 300 mV (Figure2d) that were obtained are much higher than those reported for *S. aureus* (~25 mV) and *E. coli* (~60 mV) using silicon based FET sensors.(Formisano et al. 2016; Nikkhoo et al. 2016) There are two crucial reasons for this prominent readout. First, the peptide-probes that are implemented here are small in size (~2.5 nm) and have a neutral charge that reduces the Debye screening effect and background noise, respectively. Second, the small device size (10 × 40 μm) and the measurements at the charge neutrality point enhance the sensitivity of the graphene to the charge of the bacteria. Altogether, these results confirm that G-FETs functionalized with P-KAM5_Probe are capable of detecting *S. aureus* with high specificity and sensitivity, at the single cell level. Additionally, the peptides functionalized over G-FET remain stable and were tested after storing for 24 h in PBS which showed detection capability similar to those used immediately after functionalization (Figure S4).

### 3.5. Improving sensitivity via electric field assisted bacterial binding

One potential problem associated with our design is the relatively small size of the device, which requires a high bacterial cell density (10^7^ cells/ml of *S. aureus*) to facilitate the capture of a single bacterium at the graphene surface. The active area of our G-FET is just 10 × 40 μm, while bacterial cells are distributed in an area of 2.5 × 2.5 mm, which is the size of the PDMS well placed over the device and which holds the bacterial suspension. We hypothesized that this dramatic contrast between the size of the well and that of the graphene limits the likelihood of the bacterial cells reaching the graphene surface, therefore requiring a high cell density for efficient bacterial capture. To improve the sensitivity of G-FET, we hypothesized that by applying voltage pulses from the top of the well that holds the sample, the charge of the bacteria could be exploited to drive them to the graphene surface.(Belgrader et al. 1999) Specifically, a negative voltage of −0.5 V was applied to the Pt electrode with five pulses, 10 seconds in duration to minimize potential damage to the bacteria. Figure 3a shows a clear shift in *V_D_* after electric field assisted binding at a concentration of 10^4^ cells/ml of *S. aureus*, indicating attachment of bacteria to the graphene. Increased average *V_D_* was observed with increasing concentration of cells, indicating enhanced probability of bacterial binding (see Figure 3b). However, a saturation in shift of *V_D_* was observed when the bacteria count reaches to more than 4 per device. This only occurs for concentrations above 10^6^ cells/ml, well beyond the clinically relevant limit. (Habimana et al. 2018) Moreover, electric-field assisted binding decreased the original incubation time before bacteria could be detected from 45 minutes to 5 minutes. Similar to the 45 min incubation method without electric field attachment, the Dirac voltage shift found in the electric-field assisted attachment is dependent on the number of bacteria on the device (Figure S5). Excitingly, our newly developed method of electric field assisted binding allows effective detection of *S. aureus* at 10^4^ cells/ml, which is 3 orders of magnitude lower in cell density than what is required in the absence of applying voltage pulses. Moreover, it also reduces the time needed to perform the measurements by 9-fold. To make sure that the selectivity of the devices is not affected by applying the voltage, we tested *B. subtilis* and *E. coli* using P-KAM5_probe devices, and *S. aureus* using P-KAM5_Control. No shift or bacterial attachment was observed after applying the voltage (Figure S6).

**Figure 3.**
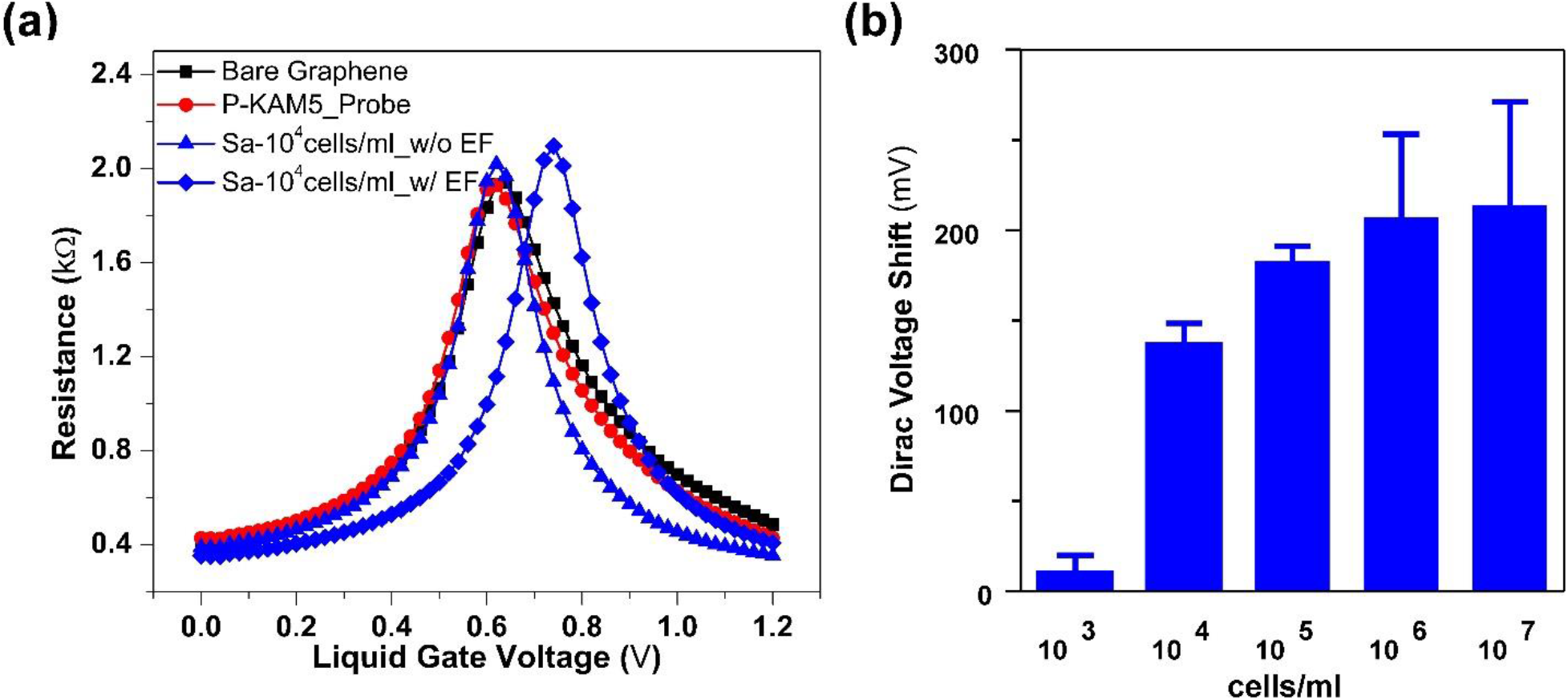
Lowering detection with electric field assisted binding. Resistance versus voltage plots of G-FET functionalized with Probe peptides (P-KAM5_Probe) (a) before (blue triangle) and after (blue diamond) electric field (EF) assisted binding of S. aureus at 10^4^ cells/ml. (b) The chart shows the average Dirac voltage shift obtained with different concentrations of S. aureus after electric field assisted binding. (Data represents average and standard deviation of at least 3 independent replicates)

### 3.6. Strain specific detection of Gram-negative antibiotic resistant pathogenic bacteria

The use of peptide probes in our G-FET design offers great versatility in terms of bacterial pathogens that can be targeted. Furthermore, as we recently demonstrated, the peptide probes can be rapidly developed to differentiate antibiotic susceptible and antibiotic-resistant strains of a bacterial pathogen. We postulated that integrating such peptide probes into G-FET would allow for specific detection of antibiotic-resistant pathogenic strains. To test this hypothesis, we turned to the peptide KAM8, which was selected to bind a colistin-resistant strain of *Acinetobacter baumannii* (AB5075 LOS−;AbR). Similar to the P-KAM5_Probe, we synthesized a pyrene conjugate of KAM8 P-KAM8_Probe as well as a control peptide (P-KAM8_Control). Functionalizing G-FET with P-KAM8_Probe or P-KAM8_Control caused no shift in *V_D_* (Figure 4a) confirming their charge neutrality. After incubation with 10^7^ cells/ml of AbR, a V_D_ shift ranging between 280-460mV was observed for devices functionalized with P-KAM8_Probe (Figure 4a) while no notable shift was measured with P-KAM8_Control (Figure S7). This shows that P-KAM8_Probe effectively captures AbR cells onto the graphene surface triggering a change in V_D_. In order to confirm the strain specificity of P-KAM8_Probe, the devices were first incubated with the non-colistin resistant wild-type strain of *A. baumannii* (AB5075; AbW) at 10^7^ cells/ml. This triggered no shift in Dirac voltage indicating that P-KAM8_Probe does not interact with the strain. Subsequently, the same device was incubated with the antibiotic-resistant strain AbR which showed a shift of 280mV confirming the probe interacting with AbR. Additionally, similar to what was observed for the interaction between *S. aureus* and P-KAM5_Probe, the measured voltage shifts correlate with the number of bacterial cells attached to the graphene surface (Figure S8). We found a single *A. baumannii* produced a *V_D_* shift of ~200mV, which is comparatively higher than that obtained with *S. aureus* which likely results from a higher density of surface charge displayed by a Gram-negative bacterium in comparison to Gram-positives.(Kłodzińska et al. 2010) Further, to see the effect of pH on bacterial binding, G-FETs functionalized with P-KAM8_Probe were tested with *A. baumannii* suspended in PBS altered to pH-6 and pH-8. Attachments were observed in both cases with variation in amount (Figure S9).

**Figure 4.**
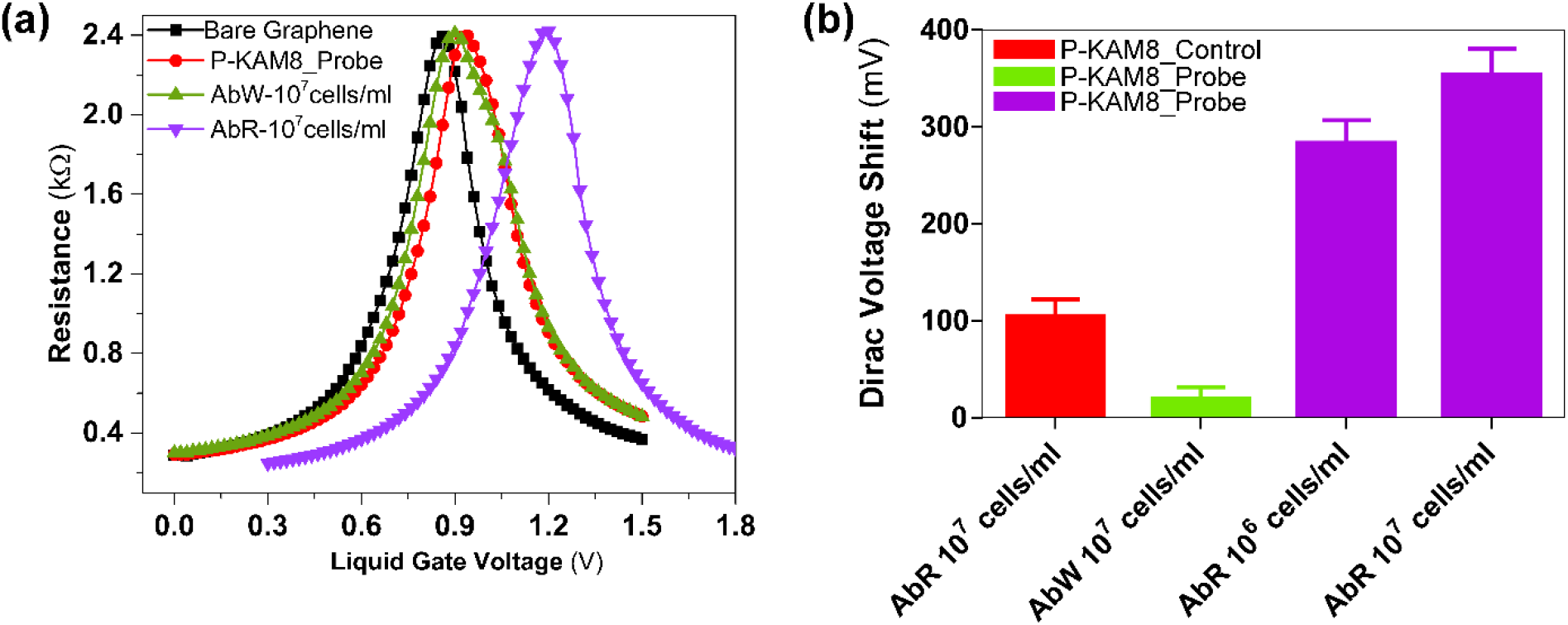
Specific detection results of A. baumannii. (a) Resistance vs voltage plots of G-FET for detection of A. baumannii with probe peptides KAM8_Probe. No shift was observed when the colistin sensitive wt A. baumannii strain (AbW) was exposed to the device, while a ~300 mV shift occurs in the presence of the colistin resistant strain AbR. (b) While P-KAM8_Control does not interact with AbR, and P-KAM_Probe does not interact with AbW, as expected only shifts in Dirac voltage are registered when P-KAM-Probe is combined with AbR. This confirms that devices functionalized with P-KAM8_Probe are specific for AbR with average voltage shifts of 280 mV and 350 mV at concentrations 10^6^ and 10^7^ cells/ml respectively. (Data represents average and standard deviation of at least 4 independent replicates)

Lastly, to determine the limit of detection of P-KAM8_Probe functionalized devices, 14 different devices were tested using suspensions of 10^7^ cells/ml and 10^6^ cells/ml of AbR, obtaining average Dirac voltage shifts of about 350 and 280 mV, respectively (Figure 4b). To reduce the required density, we again employed the electric-field assisted binding method. However, it seems that AbR requires higher voltage pulses as no shift was observed when using −0.5V, the setting that worked for *S. aureus*. Attachment of AbR was detected after applying −1V for 100s, however this seemed to damage the electrodes in the devices. We found this issue was solved by slowly sweeping the voltage from 0 to −1V with a step voltage of 10mV, resulting in detection of AbR at cell densities as low as 10^4^ cells/ml (Figure S10). Taken together, these results demonstrate the potential of the peptide-functionalized G-FETs for specific detection of antibiotic resistant strains of bacterial pathogens. Moreover, our results with at least two bacterial species (*S. aureus* and colistin resistant *A. baumannii*) suggest our new electric-field assisted binding method drastically improves the detection limit and required measurement time. The achieved lower detection limit for *S. aureus* is consistent with the recently reported values of 10^4^ CFU/ml, utilizing a long period fiber grating based biosensor and antibody probes, while better in detection time of 5 minutes as compared to 30 minutes.(Yang et al. 2019) Similarly for *A. baumannii*, the achieved detection limit is lower than the reported threshold value of 10^5^ CFU/ml as per the CDC reports.(Bulens et al. 2018)

Further reduction in the detection limit below 10^4^ cells/ml may be desired in some clinical cases which could be obtained by optimizing the device design, PDMS well size, as well as the electric field application process.(Habimana et al. 2018; Miller et al. 2018) Though, increasing the device size could increase the attachment of bacteria at lower concentrations, it may also reduce the sensitivity of the devices due to non-uniformity (e.g. wrinkles or multiple grains) and impurities with large area graphene.(Caillier et al. 2013; Mates et al. 1982) Similarly, large area devices would limit the possibility of miniaturization and multiplexing. Importantly, by applying more cycles of incubation and voltage on the same device we achieved bacterial detection even at 10^3^ cells/ml. While, these results showed more variability between replicates, it highlights that further sensitivity improvements are possible. One possible reason for the variability at 10^3^ cells/ml is that in the 20μl sample that is added to the device there are only ~20 bacterial cells present. As described above, the sensing area is small (10 × 40 μm) compared to the PDMS well that holds the 20 μl sample (2.5 × 2.5 mm). This means that the bacterial density is just 3.2/mm^2^ and thus even with the applied electric field the travel distance of a bacterium to the probes at the graphene surface remains far, with potential for other locations of attachment. Hence, optimizing the geometry of the PDMS well, resist surface and voltage application process, and/or integrating the system into a PDMS microfluidics chip could help to further reduce the detection limit. The smaller size, neutral nature (chargeless at ~pH 7) resulted in large changes in Dirac voltage (point of charge neutrality −*V_D_*) per bacterium, allowing for single cell electrical detection on G-FET. Importantly, our G-FET design enabled direct quantitative comparison of the electrical and optical readout by simple optical imaging of G-FET. The attached bacteria can also be counted by optical microscope; however, G-FETs have several advantages including easy multiplexed detection, electric field assisted binding at low concentration, user friendly operation when interfaced with electronics and can meet the requirements of point-of-care diagnostics. Electric field assisted bacterial binding overcame the tradeoff between the probability of attachment and *V_D_* shift per bacteria. The wide applicability of these peptide probes enabled detection of different pathogenic bacterial species, as well as an antibiotic resistant strain at a single cell level. Thereby our G-FET plus peptide combination offers a new promising route towards cheap, fast, multiplexed and low concentration detection of clinically relevant pathogenic bacterial species and their antibiotic resistant variants.

## 4. Conclusion

In summary, we demonstrated an electronic, label free biosensor of clinically relevant bacteria by implementing two new capabilities in G-FETs; dielectrophoresis capture of the target and highly specific synthetic peptides. This enabled smaller size (10 × 40 μm) G-FETs, which are able to selectively detect a single bacterium on the device in just 5 minutes. Furthermore this is achieved with a small (20 μl) sample volume at a detection limit of 10^4^ cells/ml, which is the threshold at which urinary tract infections (Doern and Richardson 2016; Nicolle et al. 2005) or bronchoalveolar lavage fluid are indicated to be disease causing.(Kalil et al. 2016; Miller et al. 2018) Devices showed capability for selective detection of multiple pathogenic bacterial strains including *Staphylococcus aureus* and antibiotic resistant *Acinetobacter baumannii*. Finally, the presented G-FETs design was successfully tested simultaneously for specific and unspecific peptides and can detect at least two different bacterial strains on the same platform. In the future, more detailed studies of concentration dependence are needed to achieve quantitative detection of targeted bacteria.

## Acknowledgments

N.K and K.S.B. acknowledge the primary support of the US Department of Energy (DOE), Office of Science, Office of Basic Energy Sciences under award no. DE-SC0018675. T.v.O and J.O-M were supported by R01-AI110724 and U01-AI124302. W.W. and J.G. acknowledge the National Institutes of Health financial support through grants R01-GM102735 and R01-GM124231. M.G. and acknowledge support from the National Science Foundation, through grant DMR-1709987. N.B. acknowledges support from the National Science Foundation, through grant BIO-1460628.

