## Supplementary material for "Dielectrophoresis assisted rapid, selective and single cell detection of antibiotic resistant bacteria with G-FETs": S1

**Functionalization of graphene with peptides:**

In order to achieve the uniform functionalization of peptides, we first patterned bare graphene on $SiO_2$/Si substrates so that the change in height between graphene and $SiO_2$ can be monitored. Then the patterned graphene chips were incubated with two different pyrene-conjugated peptides (P-KAM5_Probe and P-KAM8_Probe) solutions by systematically increasing the time of incubation while keeping the concentration constant. Higher concentration (10 µM) of peptides dissolved in 1x PBS was chosen to maximize/saturate the coverage of graphene surface as non-uniform coverage causes unspecific binding of bacteria. After incubation, the chips were rinsed 3 times in PBS followed by DI water to remove the peptides not attached to graphene and then dried with Argon for AFM characterization. The incubation time at which the expected increase in height obtained, was chosen to functionalize the G-FETs to perform bacteria detection. Optimum incubation time for P-KAM5_Probe and P-KAM8_Probe was found to be 2h and 16h, respectively (Figure S1a and S1b). After functionalization with peptides, an increase in height of ~2.5 nm was observed compared with the bare graphene surface.

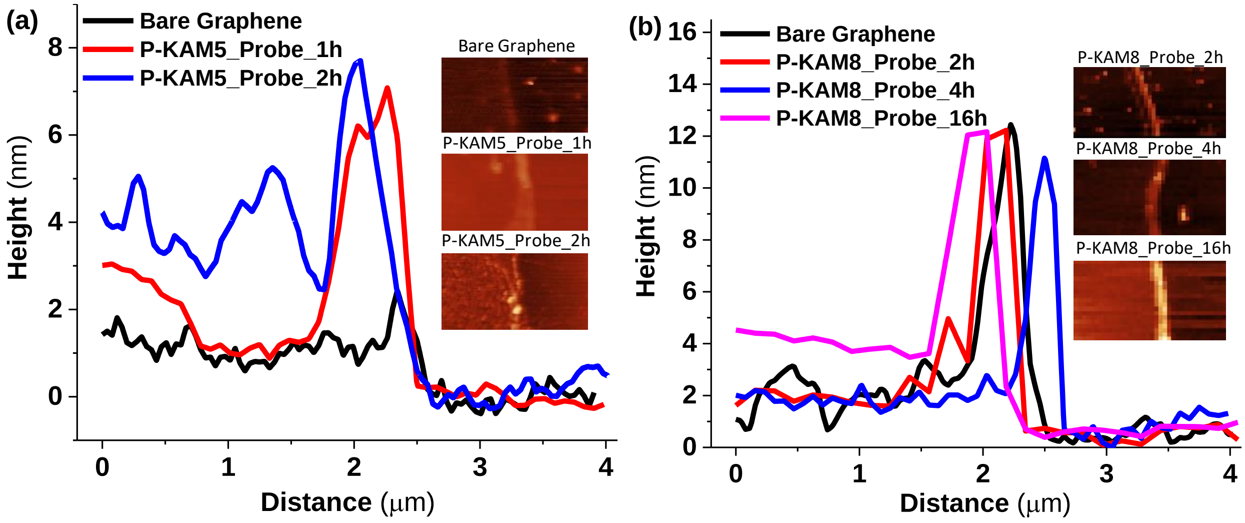

*Figure S1. The height profiles and respective AFM images of the patterned graphene before and after functionalization with peptides of 10 µM concentration at different incubation times. (a) The height profile and AFM images (inset) for bare graphene and after incubation with P-KAM5_Probe. (b) The height profile and AFM images (inset) after incubation with P-KAM8_Probe.*

**Effect of buffer strength in the measurement:**

In order to confirm the Dirac voltage shift upon bacterial binding which is known to be limited by Debye screening effect (Formisano et al. 2016; Wu et al. 2017), the G-FET was measured at two different buffer concentrations i.e. 1x PBS and 0.01x PBS having Debye length of 0.75 nm and 7.5 nm, respectively. G-FET functionalized with P-KAM5_Probe was incubated with S. aureus in 1x PBS for bacterial binding and then shift in Dirac voltage measured at two different PBS concentrations. No notable Dirac voltage shift was observed at 1x PBS while significant at 0.01x PBS, confirming the effect of Debye screening (Figure S2).

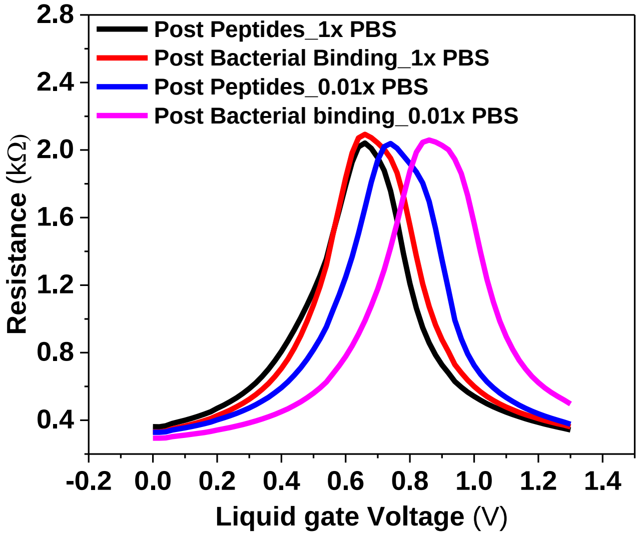


*Figure S2. G-FET characteristics before and after binding of S. aureus measured in 1x PBS and 0.01x PBS.*

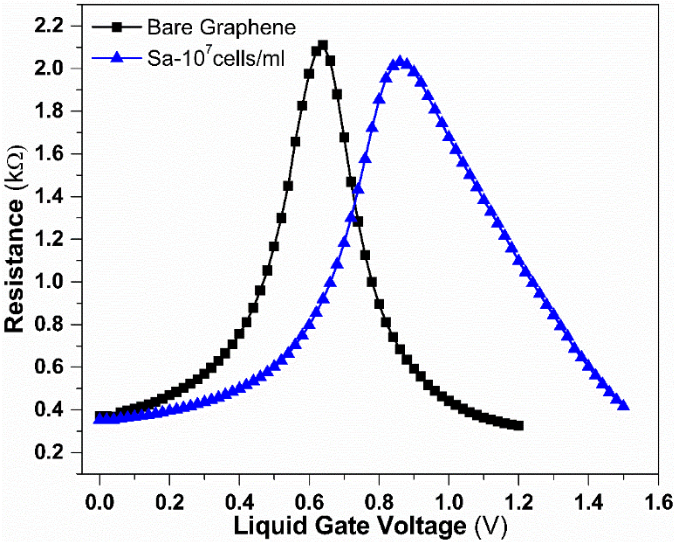


*Figure S3. G-FET characteristics before and after adsorption of S. aureus. A shift of 220mV in the Dirac voltage observed when G-FET with bare graphene was incubated with S. aureus (Blue Triangles) in comparison with the bare graphene (black squares).*

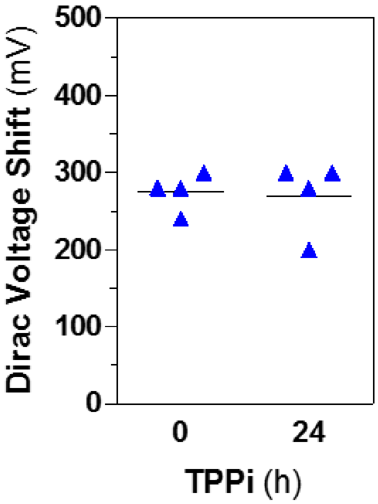


*Figure S4. Stability test, Different GFETs functionalized with P-KAM5_Probe and stored for 24h and then detection of S. aureus at $10^7$ cells/ml was performed, Measured Dirac voltage shift of different G-FETs after 0h and 24h time post probe incubation (TPPi).*

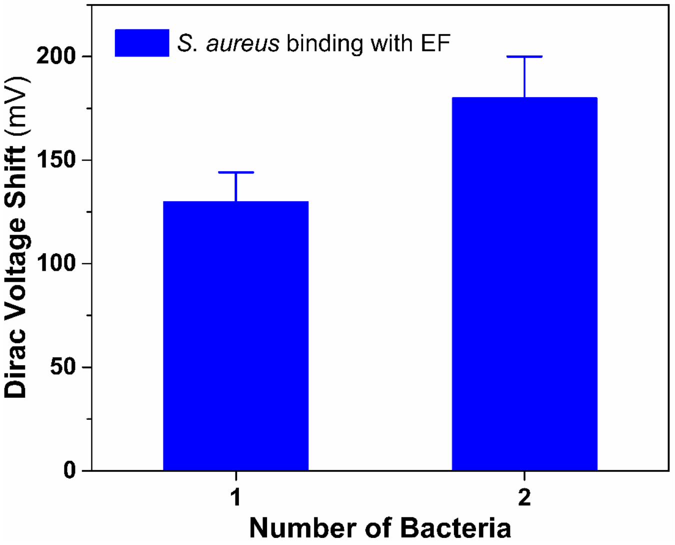


*Figure S5. Measured Dirac voltage shift of G-FETs having different number of bacteria (S. aureus) attached obtained with electric field assisted binding at a concentration of $10^4$ and $10^5$ cells/ml.*

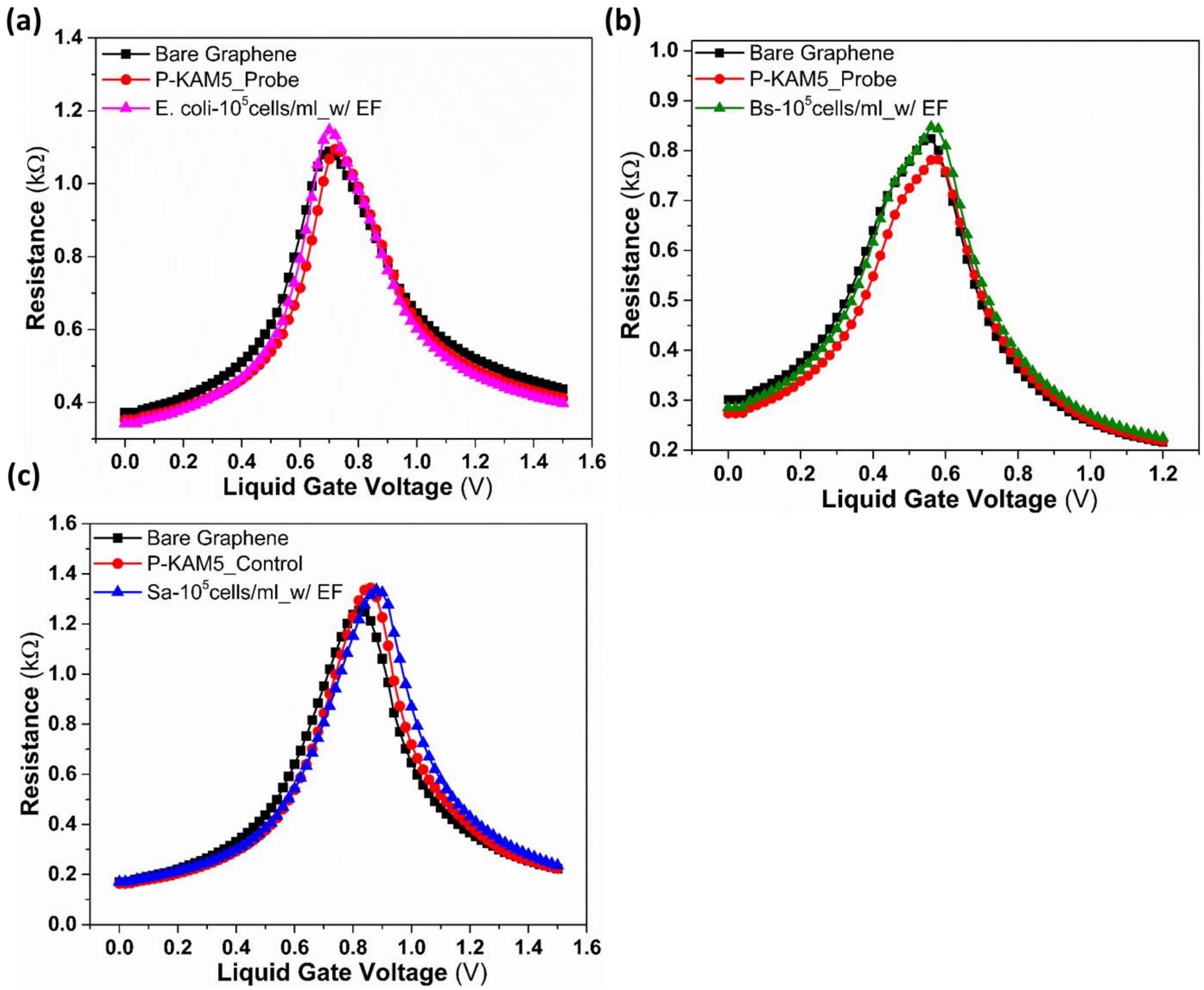


*Figure S6. Resistance versus voltage plots of G-FET functionalized with Probe peptides (P-KAM5_Probe) and after electric field assisted binding of E. coli at $10^5$cells/ml (a), B. subtilis (b). Resistance versus voltage plots of G-FET functionalized with control peptides (P-KAM5_control) and after electric field assisted binding of S. aureus at $10^5$cells/ml (c).*

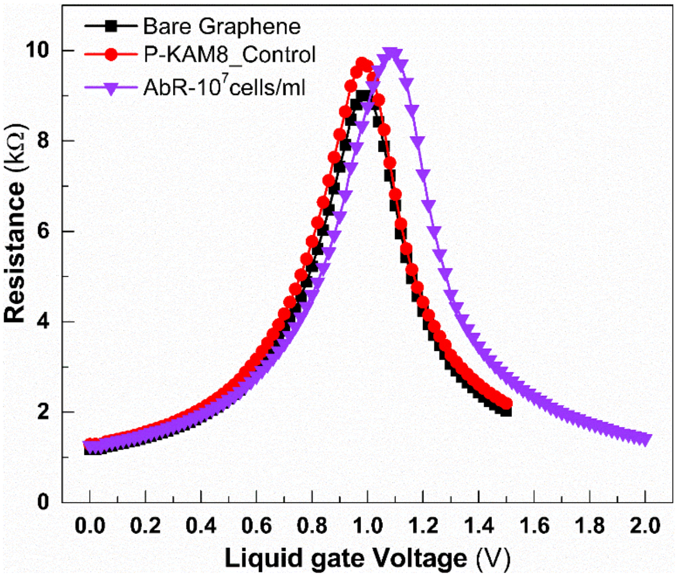


*Figure S7. Resistance vs voltage plots of G-FET for detection of S. aureus, G-FET functionalized with control peptides (P-KAM8_Control) and incubated with A. baumannii of concentration $10^7$cells/ml, no notable voltage shift observed neither with peptide and nor with bacteria and no attachment seen.*

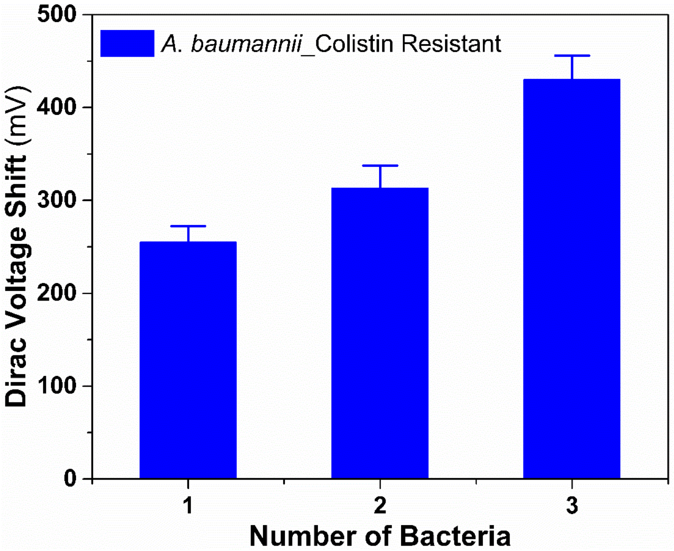


*Figure S8. Sensitivity of G-FET devices for the detection of colistin-resistant A. baumannii (AbR). Direct quantitative comparison of electrical and optical readouts of G-FETs functionalized with P-KAM8_PROBE post incubation with a $10^7$cells/ml suspension of A. baumannii AbR. Measured Dirac voltage shift of G-FETs having different number of bacteria attached, devices having single bacterium attached show an average shift of ~250mV and linear increase in voltage shift is observed with increased number of bacteria attached.*

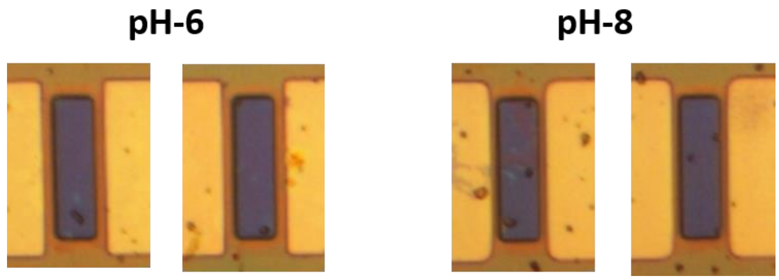


*Figure S9. Effect of pH on binding of bacteria A. baumannii. G-FETs functionalized with P-KAM8_Probe were tested with A. baumannii suspended in PBS altered to pH-6 and pH-8. Attachments were observed in both cases with variation in amount.*

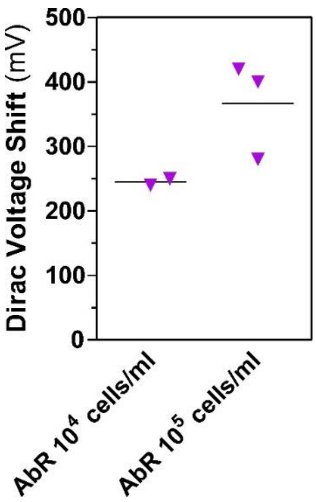


*Figure S10. Bar chart shows average Dirac voltage shift versus concentrations obtained with electric field assisted binding of bacteria A. baumannii.*
